# MIMt – A curated 16S rRNA reference database with less redundancy and higher accuracy at species-level identification

**DOI:** 10.1101/2023.12.15.571851

**Authors:** M. Pilar Cabezas, Nuno A. Fonseca, Antonio Muñoz-Mérida

**Affiliations:** Centre of Molecular and Environmental Biology (CBMA), Department of Biology, University of Minho, Campus de Gualtar, 4710-057 Braga, Portugal; Institute of Science and Innovation for Bio-Sustainability (IB-S), University of Minho, Campus de Gualtar, 4710-057 Braga, Portugal; CIBIO-InBIO, Research Center in Biodiversity and Genetic Resources, 4485-661 Vairão, Portugal; BIOPOLIS Program in Genomics, Biodiversity and Land Planning, CIBIO, Campus de Vairão, 4485-661 Vairão, Portugal

## Abstract

**Motivation:** Accurate determination and quantification of the taxonomic composition of microbial communities, especially at the species level, is one of the major issues in metagenomics. This is primarily due to the limitations of commonly used 16S rRNA reference databases, which either contain a lot of redundancy, or a high percentage of sequences with missing taxonomic information. The use of these incomplete or biased databases may lead to erroneous identifications and, thus, to erroneous conclusions regarding the ecological role and importance of those microorganisms in the ecosystem.

**Results:** The current study presents MIMt, a new 16S rRNA database for archaea and bacteria’s identification, encompassing 39 940 sequences, all precisely identified at species level. MIMt aims to be updated at least once a year to include all new sequenced species. We evaluated MIMt against Greengenes, RDP, GTDB and SILVA in terms of sequence distribution and accuracy of taxonomic assignments. Our results showed that MIMt contains less redundancy, and despite being five to 85 times smaller in size than existing databases, outperforms them in completeness and taxonomic accuracy, enabling more precise assignments at lower taxonomic ranks and thus, significantly improving species-level identification.

**Availability and Implementation:** MIMt is freely available for non-commercial purposes at https://mimt.bu.biopolis.pt

## Introduction

Microorganisms are the most diverse and abundant group of living organisms on Earth. All Bacteria, Archaea, and most lineages of the Eukarya domain belong to this group (Sattley and Madigan 2015). They live in almost every type of habitat (aquatic, terrestrial, atmospheric, or living host), and some of them thrive in extremely harsh conditions (Sattley and Madigan 2015, Poli *et al*. 2017). Although some species are harmful to certain plants and animals and may cause serious diseases in humans (e.g., tuberculosis, cholera, diphtheria, etc.), most microorganisms provide beneficial services and are essential for the proper functioning of ecosystems, playing an important role in biogeochemical cycles, aiding in the decomposition process or food fermentation, and making up the human microbiota, which is crucial for the proper functioning of the digestive and immunological systems (Vaitilingom *et al*. 2010, Sattley and Madigan 2015, Wang *et al*. 2017). However, despite their importance, most species have not been characterized so far, mainly due to the widespread lack of suitable culturing methods (Epstein 2013).

Over the past decade, the development of high-throughput sequencing technologies has expanded and altered microbiologists’ view of microorganisms and how to study them. In particular, metagenomics, the study of genetic material recovered directly from environmental samples (Ghosh 2019), has been successfully used to identify previously unrecognized species (Chalita *et al*. 2020), quantify the taxonomic composition of a microbiome (Lee *et al*. 2020; Rosado *et al*. 2023), and unravel their interactions and functions in the ecosystem (Bohan *et al*. 2017); thus, improving our knowledge and understanding of microbial diversity without prior cultivation (Handelsman 2004, Ghosh 2019).

For bacteria and archaea, the 16S small-subunit ribosomal RNA (16S rRNA) gene has been established as the ‘gold standard’ for identifying and characterizing the diversity of metagenomic samples, as it is present in almost all species, its function is highly conserved, and its size (1 550 bp) is large enough for bioinformatic purposes (Janda and Abbott 2007, Boughner and Singh 2016). The presence of hypervariable regions flanked by conserved ones, that can be targeted by universal primers, allows discrimination between different species (Janda and Abbott 2007, Boughner and Singh 2016). However, an accurate species-level classification for 16S rRNA gene sequences remains a serious challenge for microbiome researchers, because of the quality and completeness of the most used reference databases (Santamaria *et al*. 2012, Bengtsson-Palme *et al*. 2016, Brietwieser *et al*. 2019), namely Greengenes (DeSantis *et al*. 2006), the Ribosomal Database Project (RDP, Cole *et al*. 2014), SILVA (Quast *et al*. 2013) and the Genome Taxonomy Database (GTDB, Parks *et al*. 2021). These databases have several shortcomings, namely gaps in taxonomic coverage, mislabeled sequences, excessive redundancy, and incomplete annotation (especially at low taxonomic levels). In addition, these databases have different underlying taxonomies, and differ in their size, taxonomic composition, and alignment and classification approaches (Balvociute and Huson 2017, Brietwieser *et al*. 2019, Dueholm *et al*. 2020). For instance, Greengenes provides bacterial and archaeal taxonomy based on automatic *de novo* tree construction of quality-filtered sequences, but it has not been updated since 2013. Moreover, although most sequences have taxonomic information up to l family level, only over half of them are annotated at the genus level (McDonald *et al*. 2012, Parks and Won 2018). The RDP database contains bacterial and archaeal small subunit (SSU) rRNA gene sequences and fungal large subunit (LSU) rRNA gene sequences (Maidak *et al*. 2001, Cole *et al*. 2014), most of them obtained from the International Nucleotide Sequence Database Collaboration (INSDC; 27). However, this database has not been updated since September 2016. Taxonomy assignments are obtained using the Naïve Bayesian Classifier (Wang *et al*. 2007) and named according to Bergey’s taxonomy (Whitman 2015). Although most sequences have complete taxonomy up to species level, most of them are annotated as ‘uncultured’ or ‘unidentified’ taxa (Maidak *et al*. 2001, Cole *et al*. 2014). On the other hand, SILVA contains taxonomic information for all three domains of life (Pruesse *et al*. 2007, Quast *et al*. 2013, Yilmaz *et al*. 2014). It is the only database that is both manually curated and frequently updated. Taxonomic classifications follow Bergey’s taxonomy (Whitman 2015) and the List of Prokaryotic Names with Standing in Nomenclature (LPSN, Parte 2014). In the beginning, this database was designed to store all 16S rRNA sequences present in publicly available works and not to be used as a reference database for microbial identification. For this reason, its distribution was biased. However, the latest released version incorporates a non-redundant dataset (Ref NR 99) where highly identical sequences have been removed (Quast *et al*. 2013). Despite this, most sequences are identified as ‘uncultured’, thus, not providing any clue about the identity of the existing microorganisms (Park and Won 2018). Finally, GTDB was recently developed to provide a standardized bacterial and archaeal taxonomy based on genome phylogeny (Parks *et al*. 2018, 2021). It has been kept up to date till now and most sequences are identified to species level; however, it contains huge redundancy, and the taxonomic classification in some instances employs nonstandard definitions (i.e. Pseudomonas_A gstutzeri_C and Pseudomonas_A gstutzeri_H or Luteimonas gsp001717465) that inflate the database’s species counts and potentially cause errors in estimation methods (i.e., the diversity of a sample).

An accurate and reliable taxonomic identification is a critical first step in metagenomic analyses, which is highly affected by the choice of the database. The use of incomplete or biased databases, such as the previously mentioned, can lead to erroneous interpretations of the community composition; and thus, to erroneous conclusions regarding the ecological role and importance of those microorganisms in that ecosystem (Park and Won 2018, Brietwieser *et al*. 2019, Chalita *et al*. 2020). The aim of the present study was to develop a novel, compact 16S rRNA database for accurate and reliable bacteria and archaea taxonomic classification. This database excludes sequences not identified at the species level or with a vague taxonomy description, and it is intended to be used as a reference collection in any kind of microbiome studies. The performance of this new database, named MIMt (Mass Identification in Metagenomic tests), was compared against Greengenes, RDP, SILVA and GTDB databases in terms of sequence distribution and accuracy of taxonomic assignments.

## Material and methods

### MIMt construction

To construct MIMt, all representative, reference and the “latest assembly” of every genome belonging to archaea or bacteria were downloaded from the National Center for Biotechnology Information (NCBI) FTP site (ftp.ncbi.nlm.nih.gov/genomes/). When more than one genome from the same species was available, only the most recent was kept thus avoiding within-species redundancy.

For each retrieved genome, the exact location (start and end position) of 16S rRNA sequences was identified using RNAmmer 1.2 (Lagesen *et al*. 2007), which relies on Hidden Markov Models (HMMs) for both speed and accuracy. Then, the corresponding sequences were extracted and put together into a file to build the 16S MIMt database. The NCBI Taxonomy database (Federhen 2012, Schoch *et al*. 2020) was used to assign the complete taxonomy to each sequence. In this database, each taxon is identified with a stable and unique numerical identifier (taxid), which is linked to its full taxonomic classification (Federhen 2012). Taxids were obtained by entering the taxon name for each sequence in the NCBI database Taxonomy Browser and then from the NCBI taxdump file, which comprises the list of all taxid available at NCBI and their corresponding classification, it was obtained the full taxonomic information of each 16S microbial sequence. Taxonomy was subsequently formatted by adding the appropriate taxonomic rank prefixes (K_____for Kingdom, P_____ for Phylum, C_____ for Class, O_____ for Order, F_____ for Family, G_____ for Genus, and S_____ for Species), to be comparable with the most used reference databases. Sequences from uncultured or unidentified organisms and those not identified to species level were removed from the database.

### Databases to compare

The most widely used reference databases (Greengenes, RDP, SILVA and GTDB) were used to test their performance in classifying 16S sequences against MIMt. These databases are built in two standard formats: 1) including the taxonomy in the sequence file, such as RDP, SILVA and GTDB; or 2) with the taxonomy in a different file associated with the sequence ID, such as Greengenes. From the latter, the file “gg_13_8_taxonomy.txt.gz” was used. This file contains the full taxonomy for every sequence included in the database, and all taxonomy strings are strictly 7 levels and prefixed. From the RDP database, two files – one containing archaea (“current_Archaea_unaligned.fa.gz”) and other containing bacteria (“current_Bacteria_unaligned.fa.gz”) from the release 11-were downloaded and compiled into a single file. From SILVA database, the non-redundant version of the SSU Ref dataset (“SILVA_138.1_SSURef_NR99_tax_silva_trunc.fasta.gz”), recommended to be used as the reference dataset for rRNA classification (Quast *et al*. 2013), was downloaded. Finally, for the GTDB database, the file “ssu_all_r202.fna” that includes the sequence and the taxonomy itself was used.

### Species distribution in the databases

Following the taxonomic classification of every single sequence in the databases, the number of species was estimated. In addition, sequences with valid taxonomic information for each taxonomic level were counted to estimate the annotation efficiency as we move forward in the taxonomy. Thus, databases populated in sequences identified just with the environment where they were isolated will fail to classify sequences at the lower levels such as genus and species.

The redundancy of the databases was also calculated based on the number of sequences belonging to each species they contain. Considering that a bacterial genome contains from one to 15 16S rRNA molecules with an average of 4.2 molecules per genome (Větrovský 2013), it is expected to find some redundancy just because of the nature of the sequence itself. Nevertheless, any number higher than that will indicate that the database contains repeated sequences from different isolates or cultures that do not provide extra information and reduce efficiency in classifying species. To verify this bias, all sequences belonging to the same species were grouped and the species from highest to lowest representation were ordered. Then, the ten most represented species of the set were taken and the percentage that represents over the total was calculated.

The list of species present in each database was used to evaluate the similarity between the different databases through a Venn diagram using the library VennDiagram (Chen and Boutros 2011) in RStudio v. 2022.12.0+353 (RStudio Team 2020). The species shared by all or some of the databases will give an idea of the specificity of each database when assigning the taxonomy.

### Classification performance test

To assess MIMt’s performance against existing databases we replicated the analyses of two previously published studies (Estrella-González *et al*. 2020, Lee *et al*. 2020) based on two completely different scenarios. The first included samples from colon, cloaca, and magnum from 34-week-old laying chickens (accession number: PRJNA604381), and the second focused on microorganism biodiversity in compost from different sources (accession number: MG-RAST mgp94523). Only one replicate per condition was used: 1) Cloaca sample (SRR10998526), magnum sample (SRR10998535), and colon sample (SRR10998554) from the chicken gut microbiota study; and 2) Vegetable waste (sample 3 rep.1), “alpeorujo” (sample 2 rep. 1), sewage sludge (sample 3 rep.2), agro waste (sample 1 rep. 3) and urban solid waste (sample 2 rep. 3) from the compost microbiota study.

These datasets were processed using three of the most popular algorithms in metagenomic analysis: QIIME2 (Bolyen *et al*. 2019); DADA2 (Callahan *et al*. 2016); and Mothur (Schloss *et al*. 2009). When using QIIME2, sequences were denoised using DADA2, quality filtered, clustered into Operational taxonomic units (OTUs), and subsequently assigned to a reference taxonomy. In the case of DADA2, the protocol consisted of filtering and trimming of the sequences, learning the error rates, sample inference, merging of paired reads (not applicable to the chicken study for being single-end sequencing), and taxonomy assignment of the Amplicon Sequence Variants (ASVs) obtained. Finally, the protocol followed when using Mothur consisted of transforming the fastq files to fasta and then assigning the taxonomy to each of the sequences using the function classify.seqs. All the analyses were carried out on a server with 64 cores and 256 GB of RAM.

The analysis of the results considered the sequences annotated using each database, the number of distinct species identified, and the time spent on taxonomic identification. In some cases, the taxonomic assignment was performed on OTUs or ASVs, while in the case of Mothur it was performed directly on each of the sequences of the original dataset. For that reason, the results had to be normalized to the final number of sequences to be annotated after the preprocessing.

For the final count of classified sequences all values showing simply a vague description of the sequence or internal codes were excluded (i.e. marine bacterium, unclassified bacteriales, uncultured archaeon, and sp22811).

## Results

### Database composition

To construct the new 16S reference database for microorganism identification, a total of 18 317 genomes, 692 belonging to archaea and 17 625 from bacteria, were downloaded from NCBI. A total of 39 940 16S rRNA molecules were identified along the genomes which became the MIMt database (Table 1). The number of 16S molecules across species varies between 1 and 37 in *Tumebacillus avium*, with an average of 2.18 which indicates that MIMt lacks redundancy.

**Table 1.** Number of sequences and unique species at the different databases.

| Database | Sequences | Unique species | Bacteria Species | Archaea Species |
| --- | --- | --- | --- | --- |
| MIMt | 39 940 | 18 318 | 17 626 | 692 |
| GTDB | 486 035 | 12 684 | 12 217 | 467 |
| Greengenes | 203 452 | 3 032 | 2 923 | 109 |
| RDP | 3 356 809 | 11 999 | 11 501 | 498 |
| SILVA | 510 509 | 15 644 | 15 110 | 534 |

Every single sequence that constitutes the MIMt database was identified to species level (Figure 1A) throughout the NCBI Taxonomy. Moreover, most sequences have taxonomic information at all classification levels (Figure 1A), being class and family the ones with the lowest percentage of sequences assigned (99.25% and 98.51%, respectively).

**Figure 1.**
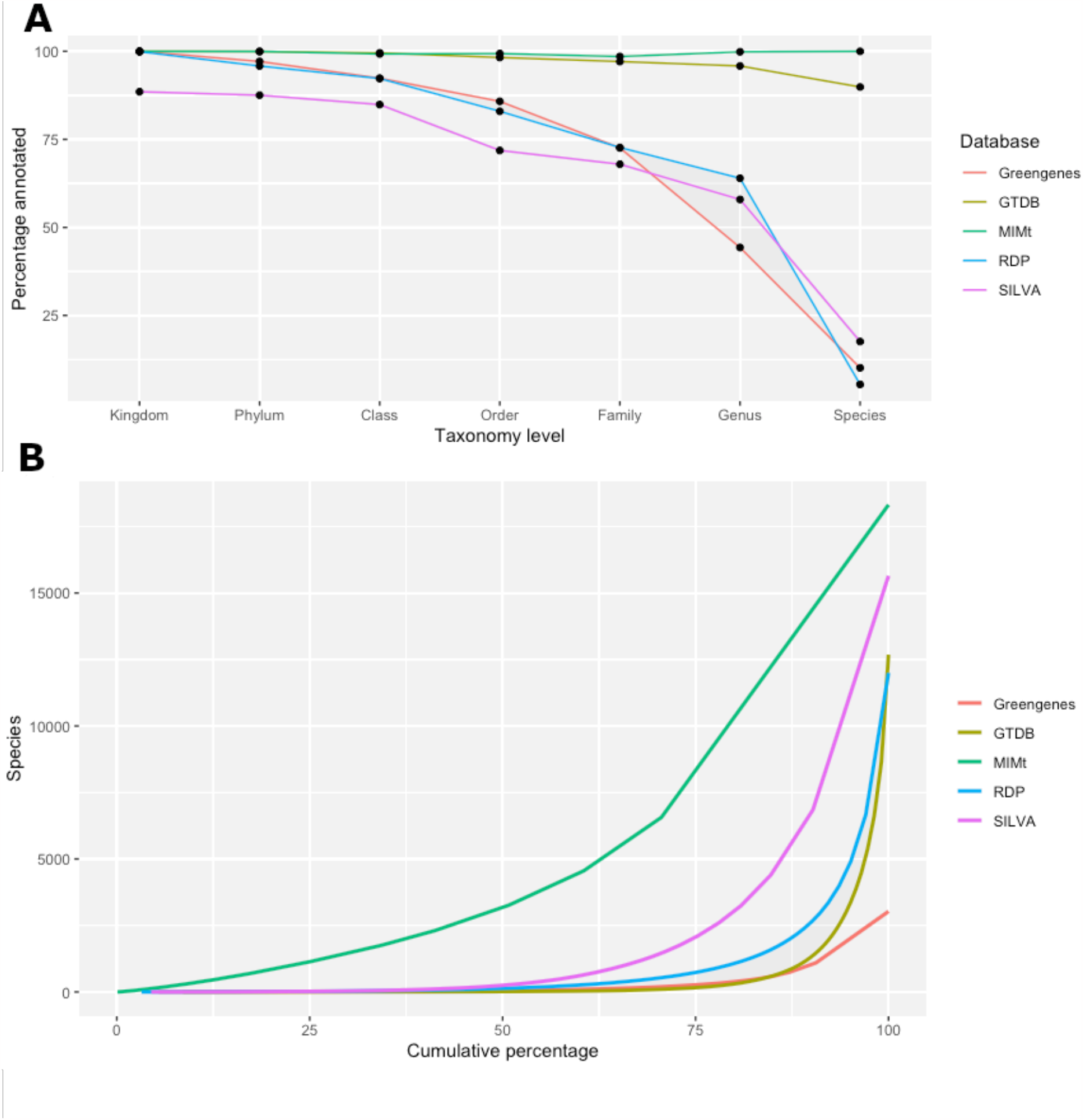
A) Distribution of sequences with meaningful information at every taxonomic level. B) Distribution of sequences per species at different databases (Greengenes, GTDB, MIMt, RDP and SILVA) expressed in cumulative percentage.

Our results show that the other databases are also consistent at higher taxonomic levels (kingdom, phylum, class and order), with around 80% of the sequences taxonomically annotated (Figure 1A). However, in Greengenes, RDP and SILVA, less than 60% of the sequences have valid taxonomic information at genus-level, and the percentage drops below 20% at the species-level (Figure 1A). GTDB is the only database that maintains acceptable levels of sequences with valid taxonomy at genus and species levels with almost 96% and 90% of sequences annotated, respectively (Figure 1A; Table 1 Supplementary Material). It is worth mentioning that RDP and SILVA databases contain most of their sequences annotated at species level, however most of them contain terms like “Taxon”, “genomosp.”, “endosymbiont”, thus being considered as uninformative. Greengenes has a low proportion of sequences taxonomically annotated up to species level (∼10%). However, it exclusively contains valid species names, therefore does not contain uninformative species names.

On the other hand, the analyses of sequence distribution at species level, revealed that MIMt, as expected, contains minimal redundancy resulting directly from the very abundance of 16S molecules in each genome (Figure 1B, Table 2 Supplementary Material). Our results showed that GTDB and Greengenes are the databases with the highest redundancy (Figure 1B). These two databases showed a similar tendency reaching 75% of the sequences annotated at species level with only 164 and 270 unique species respectively. *Escherichia flexneri* was the most represented species in GTDB with 37 521 sequences and *Faecalibacterium prausnitzii* the most frequent one in Greengenes with 1 114 sequences (percentages in the graph were relativized to the total sequences annotated at species level). The RDP database contains a bit less redundancy (Figure 1B) but still 75% of the whole database sequences belong to 735 species out of the 11 999 that it contains. Finally, for the SILVA database 75% of the sequences belong to 2 065 species from the 15 644 it has in total.

Another important aspect to analyze is the representation of species in the different databases to see if all share the same species or if there are exclusive species just in any of them. Regarding this, our results showed that only 1 465 species are common between all databases (Figure 2). Each database has a variable number of unique species, i.e. not shared with the rest, being this number significantly greater in MIMt. A fairly large number of species can also be observed shared between all databases except Greengenes (5 513) due to the smaller number of species contained in this database (Figure 2).

**Figure 2.**
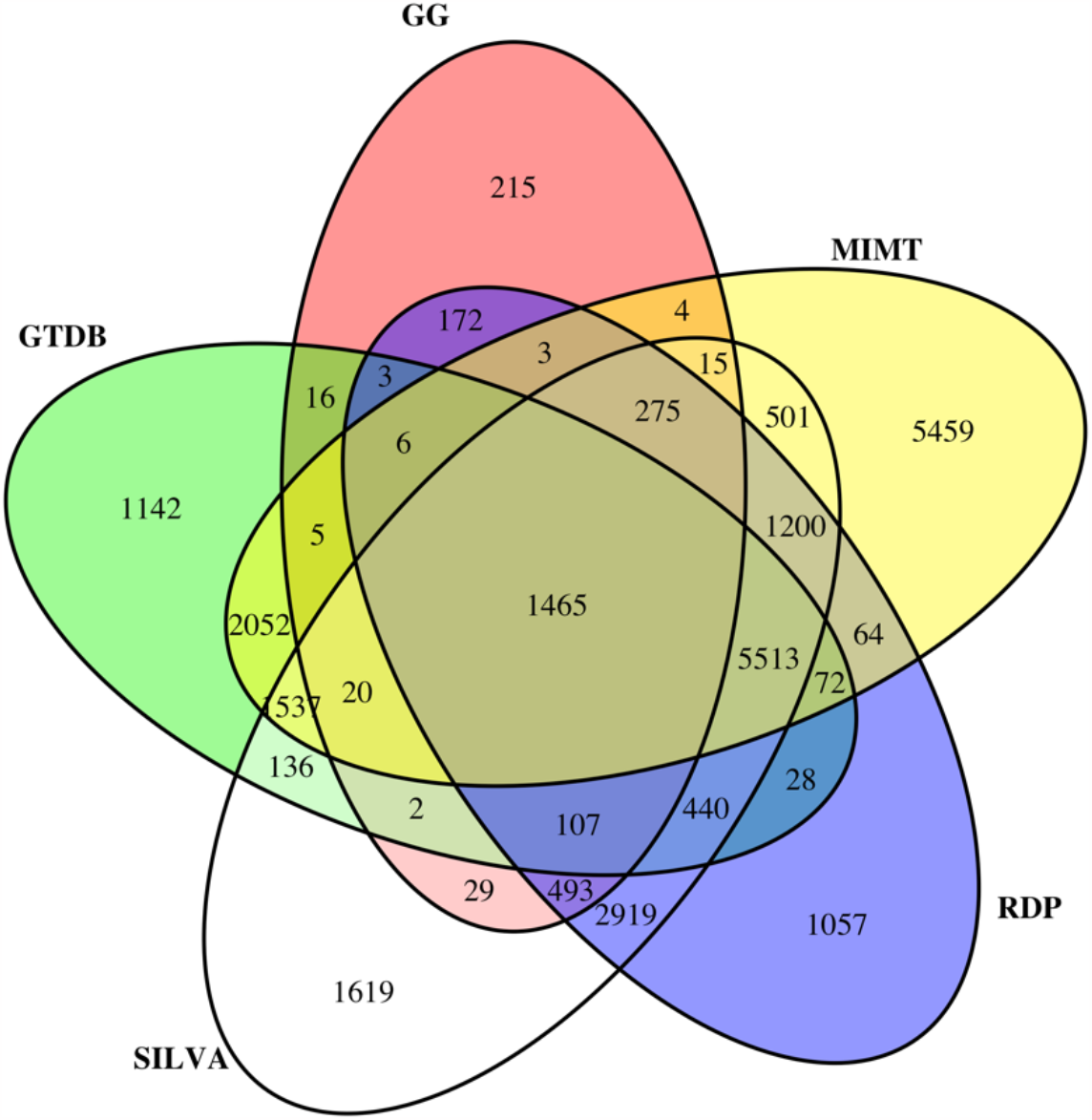
Venn diagram of the species shared between databases: MIMt, Greengenes (GG), the Ribosomal Database Project (RDP), SILVA and the Genome Taxonomy Database (GTDB).

### Taxonomic classification test

To establish the actual taxonomic allocation capacity of MIMt, 16S rRNA data from two studies already published were analyzed using QIIME2, DADA2 and Mothur. The datasets include data from different sources such as poultry intestine, differentiating between cloaca (A), magnum (B), and colon (C); and compost microbiota from vegetable waste (D), “alpeorujo” (E), sewage sludge (F), agricultural waste (G) and urban solid waste (H). In each case, the time it took to classify the datasets, the number of sequences annotated at the species level and the number of different species identified were measured (Figure 3).

**Figure 3.**
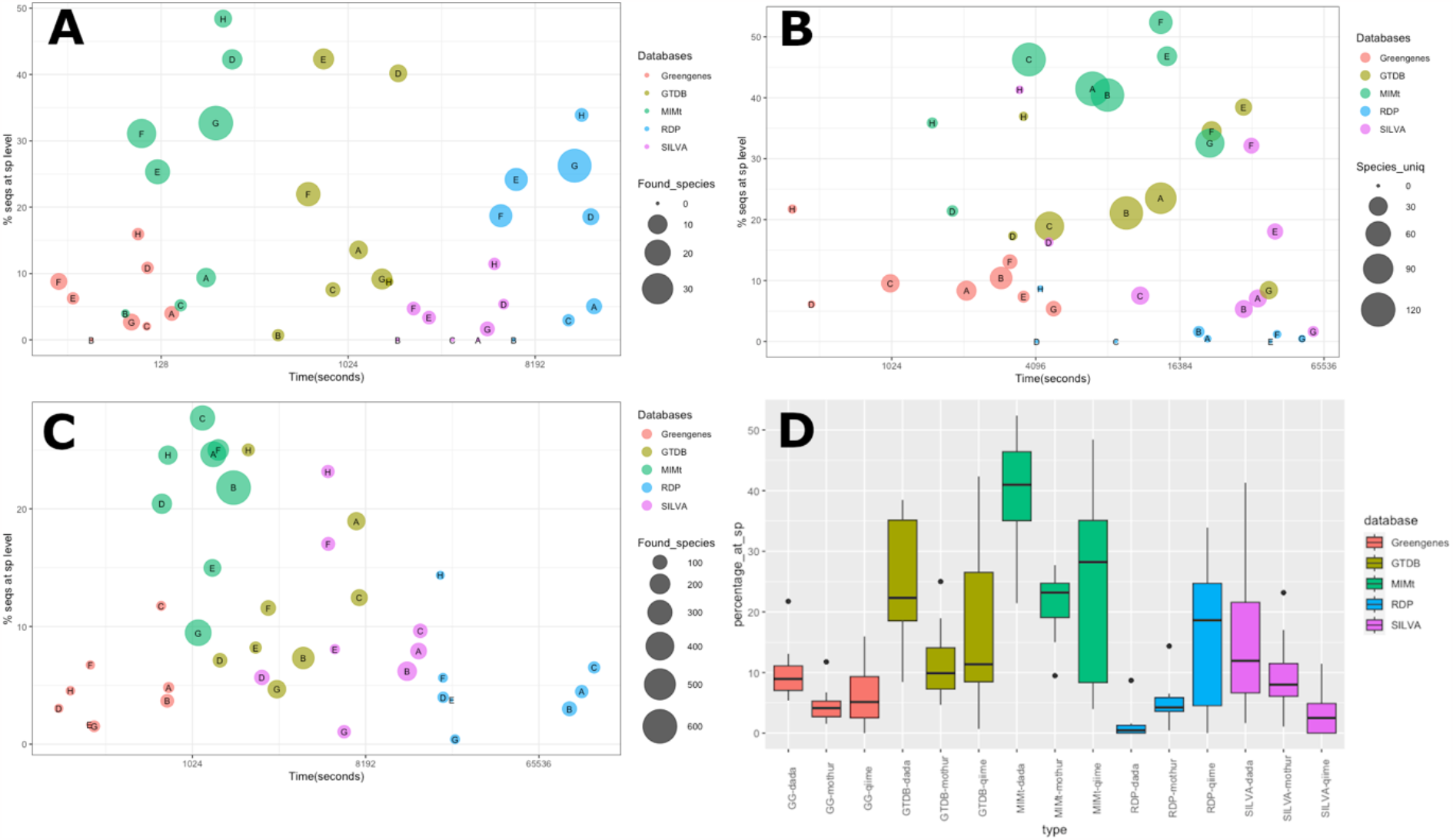
Performance evaluation in sequence classification by each database using (A) QIIME2, (B) DADA2, and (C) Mothur. Plots represent the percentage of sequences classified at species level and the time spent on the classification. The size of de bubbles represents the number of different species identified. (D) Boxplot of the classified sequences at species level in each condition.

The results show that the use of Greengenes as a database was in all cases the fastest scenario, followed by MIMt, GTDB, SILVA and RDP (Figures 3A-3C). The time spent on the analyses exhibited to be proportional to the size of the database and to the annotation structure of each database. For instance, GTDB and SILVA have a similar number of sequences, but since GTDB sequences are better annotated at all levels, resulting in a higher efficiency in the time spent in classification. MIMt showed an exceptional ratio between the time spent on classifying the datasets and the percentage of sequences annotated at the species level becoming the database with the largest number of sequences classified and the highest number of species identified (Figure 3; Table 3-5 Supplementary Material).

## Discussion

MIMt is a compact, non-redundant 16S rRNA database with all its sequences annotated to the species level. Despite its smaller size compared to other databases, MIMt has demonstrated a very good performance in metagenomic sequences classification. In the case studies considered, MIMt successfully classified over 50% more sequences than any other database (Figure 3D). The taxonomic classification tests carried out in this study show MIMt’s good performance in all conditions, making it particularly helpful in the characterization at the species level of a metagenomic sample. This is particularly useful in detection of pathogenic microorganisms or in diversity tests (alpha or beta) where incomplete or ambiguous descriptions in the taxonomy can lead to inaccurate conclusions. When is needed to identify a pathogenic microorganism in a sample or, on the contrary, beneficial agents as happens in fecal transplants, it is of great importance that apart from finding a good result against some of the sequences of the database, that sequence should be annotated accordingly or else uncertainty about the nature of the micro-organism would rule out the result.

Each sequence in MIMt represents a complete 16S molecule, that encompasses both variable and constant regions. It includes only molecules from each representative species what ensures that sequencing of any part of the molecule can be covered while avoiding having redundant sequences, which entails a substantial decrease in the time required to classify a dataset.

It should be highlighted that MIMt contains more than 5,000 species of bacteria and archaea that are not present in any of the other databases analysed (Figure 2). Furthermore, every single sequence in MIMt comes from a sequenced organism that was identified and deposited in the NCBI database. This approach avoids the inclusion of isolated fragments corresponding to 16S molecules that may be taxonomically misallocated due to contamination.

When taxonomically classifying a sequence, it is easy to perceive that we will get better results when the database contains a sequence that although less similar to our query, is well annotated at all levels, rather than if we have exactly the same sequence but its annotation is vague and incomplete. Traditionally there is a great effort for databases just to grow without putting too much emphasis on an aspect as crucial as quality. When a database used to assign taxonomy grows in number of sequences but these lack information at low levels or such information is merely descriptive without providing real taxonomy, when we compare our dataset against it, the search time will increase logically due to the size increase, but as drawback, the only benefit we will have is to classify some more sequences with just a vague description, avoiding that sequence to be annotated with another sequence of the database that, although similar to a lesser degree, does have taxonomic information. As the number of sequences in the database increases, it is easier to find cross-species identities so no taxonomic assignment will be performed due to ambiguity. It will increase the probability of finding the same sequence or one of the same species in the database but the probability of finding a cross match increases significantly more resulting in a loss in classification efficiency.

Based on the good results demonstrated by MIMt in this work, we present this new database as a powerful tool in the identification of metagenomic samples that consumes very few computational resources for a fast and efficient taxonomic classification.

## Acknowledgements

Work co-funded by the project NORTE-01-0246-FEDER-000063, supported by Norte Portugal Regional Operational Programme (NORTE2020), under the PORTUGAL 2020 Partnership Agreement, through the European Regional Development Fund (ERDF).

## References

Balvociute M, Huson DH. SILVA, RDP, Greengenes, NCBI and OTT - how do these taxonomies compare? BMC Genom 2017; 18(Suppl 2): 114.

Bengtsson-Palme J et al. Strategies to improve usability and preserve accuracy in biological sequence databases. Proteomics 2016; 16: 2454–60.

Bohan DA et al. Next-Generation Global Biomonitoring: Large-scale, Automated Reconstruction of Ecological Networks. Trends Ecol Evo 2017; 32: 477–87.

Bolyen E et al. Reproducible, interactive, scalable and extensible microbiome data science using QIIME 2. Nat Biotechnol 2019; 37: 852–7.

Brietwieser FP, Lu J, Salzber SL. A review of methods and databases for metagenomic classification and assembly. Brief Bioinformatics 2019; 20: 1125–39.

Boughner LA, Singh P. Microbial Ecology: where are we now? Postdoc J 2016; 4: 3e17.

Callahan BJ et al. DADA2: High-resolution sample inference from Illumina amplicon data. Nat Methods 2016; 13(7): 581–3.

Chalita M et al. Improved Metagenomic Taxonomic Profiling Using a Curated Core Gene-Based Bacterial Database Reveals Unrecognized Species in the Genus Streptococcus. Pathogens 2020; 9: 204.

Chen H, Boutros PC. VennDiagram: a package for the generation of highly-customizable Venn and Euler diagrams in R. BMC Bioinformatics 2011; 12: 35.

Cole JR et al. Ribosomal database project: data and tools for highthroughput rRNA analysis. Nucleic Acids Res 2014; 42: D633–42.

DeSantis TZ et al. Greengenes, a chimera-checked 16S rRNA gene database and workbench compatible with ARB. Appl Environ Microbiol 2006; 72: 5069–72.

Dueholm MS et al. Generation of comprehensive ecosystems-specific reference databases with species-level resolution by high-throughput full-length 16S rRNA gene sequencing and automated taxonomy assignment (AutoTax). mBio 2020; 11(5): e01557–20.

Epstein SS. The phenomenon of microbial uncultivability. Curr Opin Microbiol 2013; 16: 636–42.

Estrella-González MJ et al. Uncovering new indicators to predict stability, maturity and biodiversity of compost on an industrial scale. Bioresour Technol 2020; 313: 123557.

Federhen S. The NCBI taxonomy database. Nucleic Acids Res 2012; 40: D136–43.

Ghosh A, Mehta A, Khan AM. Metagenomic Analysis and its Applications. J Bioinform Comput Biol 2019; 3: 184–93.

Handelsman J. Metagenomics: application of genomics to uncultured microorganisms. Microbiol Mol Biol Rev 2004; 68: 669–85.

Janda JM, Abbott SL. 16S rRNA Gene Sequencing for Bacterial Identification in the Diagnostic Laboratory: Pluses, Perils, and Pitfalls. J Clin Microbiol 2007; 45: 2761–64.

Lagesen K et al. (2007) RNAmmer: consistent and rapid annotation of ribosomal RNA genes. Nucleic Acids Res 2007; 35: 3100–8.

Lee SJ et al. Comparison of microbiota in the cloaca, colon, and magnum of layer chicken. PLoS ONE 2020; 15(8): e0237108.

Maidak BL et al. The RDP-II (Ribosomal Database Project). Nucleic Acids Res 2001; 29: 173–4.

McDonald D et al. An improved greengenes taxonomy with explicit ranks for ecological and evolutionary analyses of bacteria and archaea. ISME J 2012; 6: 610–8.

Park SC, Won S. Evaluation of 16S rRNA Databases for Taxonomic Assignments Using Mock Community. Genomics Inform 2018; 16: e24.

Parks DH et al. A standardized bacterial taxonomy based on genome phylogeny substantially revises the tree of life. Nat Biotechnol 2018; 36: 996–1004.

Parks DH et al. GTDB: an ongoing census of bacterial and archaeal diversity through a phylogenetically consistent, rank normalized and complete genome-based taxonomy. Nucleic Acids Res 2021; 50(Database issue): D785–94.

Parte AC (2014). LPSN—list of prokaryotic names with standing in nomenclature. Nucleic Acids Res 2014; 42 (Database issue): 613–6.

Poli A et al. Microbial Diversity in Extreme Marine Habitats and Their Biomolecules. Microorganisms 2017; 5(2): 25.

Pruesse E et al. SILVA: a comprehensive online resource for quality checked and aligned ribosomal RNA sequence data compatible with ARB. Nucleic Acids Res 2007; 35: 7188–96.

Quast C et al. The SILVA ribosomal RNA gene database project: improved data processing and web-based tools. Nucleic Acids Res 2013; 41: D590–96.

Rosado D et al. Disruption of the skin, gill, and gut mucosae microbiome of gilthead seabream fingerlings after bacterial infection and antibiotic treatment. FEMS Microbes 2023; 4: xtad011.

RStudio Team (2020). RStudio: Integrated Development for R. RStudio, PBC, Boston, MA. http://www.rstudio.com/.

Santamaria M et al. Reference databases for taxonomic assignment in metagenomics. Brief Bioinform 2012; 13: 682–95.

Sattley WM, Madigan MT. Microbiology. eLS 2015; 1–10.

Schloss PD et al. Introducing mothur: open-source, platform-independent, community-supported software for describing and comparing microbial communities. Appl Environ Microbiol 2009; 75: 7537–41.

Schoch CL et al. NCBI Taxonomy: a comprehensive update on curation, resources and tools. Database 2020; baaa062.

Vaitilingom M et al. Contribution of microbial activity to carbon chemistry in clouds. Appl Environ Microbiol 2010; 76: 23–9.

Větrovský T, Baldrian P. The Variability of the 16S rRNA Gene in Bacterial Genomes and Its Consequences for Bacterial Community Analyses. PLoS One 2013; 8(2): e57923.

Wang B et al. The human microbiota in health and disease. Engineering 2017; 3: 71–82.

Wang QG et al. Naïve Bayesian Classifier for Rapid Assignment of rRNA Sequences into the New Bacterial Taxonomy. Appl Environ Microbiol 2007; 73: 5261–7.

Whitman WB. Bergey’s Manual of Systematic of Archaea and Bacteria. New Jersey, EUA: Wiley Online Library, 2015.

Yilmaz P et al. The SILVA and “All-species Living Tree Project (LTP)” taxonomic frameworks. Nucleic Acids Res 2014; 42: D643–8.

